# Birth Order Differences in Education Originate in Post-Natal Environments

**DOI:** 10.1101/2021.06.01.446519

**Authors:** Martin Arstad Isungset, Jeremy Freese, Ole A. Andreassen, Torkild Hovde Lyngstad

**Affiliations:** Department of Sociology and Human Geography, University of Oslo, PO Box 1096 Blindern, 0317 Oslo, Norway; Department of Sociology, Stanford University, Stanford, CA 94305, United States of America; Institute of Clinical Medicine, University of Oslo, PO box 4956 Nydalen, 0424 Oslo, Norway; NORMENT, Division of Mental Health and Addiction, Oslo University Hospital, Oslo, Norway

## Abstract

Siblings share many environments and much of their genetics. Yet, siblings turn out different. Intelligence and education are influenced by birth order, with earlier-born siblings outperforming later-borns. We investigate whether birth order differences in education are caused by biological differences present at birth, that is, genetic differences or in-utero differences. Using family data that spans two generations, combining registry, survey, and genotype information, this study is based on the Norwegian Mother, Father and Child Cohort Study (MoBa). We show that there are no genetic differences by birth order as captured by polygenic scores (PGSs) for educational attainment. Furthermore, we show that earlier-born have lower birth weight than later-born, indicating worse uterine environments. Educational outcomes are still higher for earlier-born children when we adjust for PGSs and in utero variables, indicating that birth order differences arise post-natally. Finally, we consider potential environmental influences, such as differences according to maternal age, parental educational attainment, and sibling genetic nurture. We show that birth order differences are not biological in origin, but pinning down their specific causes remains elusive.

## Introduction

Sibling differences receive less attention than sibling similarities, although they make up an important part of the picture of social inequality. Even for socio-economic outcomes, inequality within families (i.e., among adult siblings) can be as large as inequality between families (1). While evidence indicates a large role for social environmental influences in producing differences among siblings in the same family, documenting specific, systematic influences has been more elusive. Developmental scientists have talked about a “gloomy prospect” in which nearly all influence in and by social environments is idiosyncratic (2, 3).

Birth order has long been offered as an example of a systematic source of environmental differentiation within families. Empirically, firstborn siblings have slightly higher intelligence (4–6), educational achievement (7, 8), and income (9, 10) than their siblings born later. These differences are routinely interpreted as reflecting causal mechanisms related to childhood experiences. For example, some work suggests that parental resources are more diluted for later-born siblings as a result of competing demands for parental attention (11, 12), while other work considers whether the presence of older siblings adversely affects the cognitive environment in which younger siblings are raised (13, 14).

Even though birth order differences in achievements are routinely presumed to reflect different social environments between siblings, there are several reasons to worry they may be caused biologically.

For one, biological differences may be induced by fertility decision-making. Genetic differences among siblings have been described as a “lottery” (15–17), and parents may be more likely to have an additional child if their already-born children evince desirable traits consistent with a favorable draw from this “lottery” (18). Later-born children may exhibit a “regression to the mean” phenomenon of less propitious genetic endowments for these same traits. When deciding whether to have another child, parents have information on behavior and traits of their current child(ren) that are correlated with genotype. Predictive information about a child’s later cognitive ability is measurable at least as early as 10 months (19). Consistent with the findings of another study (20), in our data, we find that the polygenic score for educational attainment is inversely associated with a number of behaviors at age 2-3, including some net of parent’s polygenic scores (Supplementary Figure S1).

If parents adjust their fertility-behavior dependent on the first child’s behaviors in childhood and those behaviors are partly influenced by genetics, this could lead to differences in genetics between siblings according to birth order. Studies show that parents adjust their parenting according to child genotype too, engaging in more cognitively stimulating activities with children with higher PGS for education (21, 22). Thus gene-environment correlations could exacerbate any genetic or other biological differences present at birth.

Biological differences could also result from involuntary processes. Miscarriage occurs in 8-30% of all pregnancies. The risk of miscarriage increases with maternal age, but also, net of mother’s age, studies have found that number of previous pregnancies is associated with likelihood of miscarriage, albeit not in a consistent direction (23, 24). Most miscarriages happen early in the pregnancy, with the main factors being genetic abnormalities (i.e. chromosomal aberrations like autosomal trisomy) and uterine malformation (25). Changing risk of miscarriage by either factor over pregnancies, or their combination, could result in survivorship-based biological differences by birth order in live births.

Maternal age also may influence in-utero environments (26, 27), such as the rate of antibody attacks (28). Beyond this, maternal nutrition, stress, medical professional visits, and other health-related behaviors may vary according to birth order (29), causing differences in birth weight and other outcomes which influence intelligence and educational attainments later in life (30, 31). Other conditions, such as preeclampsia, have been linked to birth order (32), health, and cognition, although often not in ways consistent with the expectation of firstborn advantage (33, 34).

For that matter, mutations also rise with parental age, in particular with paternal age at conception (26). There is evidence that advanced paternal age increases the probability of offspring intelligence disorder, schizophrenia, and autism (35–37), as well as the rate of several other health-related traits (38). A consequence of these many unknowns may be that there are biological mechanisms that produce differences between siblings present at birth which we are still learning about. These can result in biological differences by birth order, albeit not necessarily a causal effect of birth order as such.

Against this backdrop it is important to recognize the limited research designs on which a considerable portion of the existing literature on birth order and educational outcomes is based. Many studies do not compare siblings in the same family, leaving genetic differences across families as an uncontrolled confounder (6). Birth-order studies often define birth order by the rearing family or by siblings who share a mother (7, 39), so that genetic differences by birth order may result from paternal genetic differences by siblings who do not share a father. Even when studies intend to include only full siblings, this is usually based only on maternal self-report. Sometimes mothers are unsure themselves who the father is (40) or misreport paternity for other reasons. For half-siblings with different fathers, paternal genetic differences could be associated with birth order, as mothers who have children with multiple partners typically have children with less educated fathers for their later-born children (41, 42).

Our study assesses birth order differences in educational outcomes within families using genetically-confirmed full siblings. We do so with data that combines registry, survey, and genotype information from families in Norway (see Methods). At stake is that, if birth order differences are not due to post-birth environments, this would imply that using birth order as an exemplar of a documented systematic effect of family environments on sibling differentiation is incorrect, furthering contentions that intrafamilial environments are far more inert than many suppose. Moreover, if differences were specifically genetic, it would also undermine various causal inference strategies that effectively assume genetic differences among full siblings are independent (18, 43). We examine whether birth order effects on education are influenced by genetic differences as measured by polygenic scores for educational attainment. We also consider broader pre-birth biological differences by looking at whether there are birth order differences in birth weight and birth length, as indicators of a combination of genetic and *in utero* factors. Furthermore, we look at non-transmitted alleles of parents, whose influence on development is sometimes referred to as “genetic nurture.”

In all, we address three research questions: 1) Is there a difference in genetics associated with educational attainment between birth orders? 2) Are there differences in birth weight and birth length between birth orders? 3) Does the putatively socially-based effect of birth order remain after accounting for genetic differences and in-utero variables? After having established that birth order differences are not biological in origin, we investigate further what may cause these differences, considering whether birth order differences vary by polygenic scores, family background, and non-transmitted parental- and sibling alleles.

## Results

We begin by analyzing the full population of Norway using administrative data to see whether there are birth order differences among children and adults. To better parallel subsequent analyses with genomic data, we exclude participants whose parents were not born in Norway (see Methods). For the population of children we study (b. 1994-2009), we examined performance on national tests by computing the child’s mean score of tests in three subjects measured at three times (grades 5, 8, and 9; standardized within subject, age, and cohort; N=301,795). For the adult population (b. 1945-1988), our outcome is completed years of education at age 30 (standardized within cohort, N=2,067,878). We use family-level fixed effects when estimating the bivariate association between birth order and these outcomes, meaning we are comparing siblings within the same family. We control for sex and maternal age (44), and run the models separately by sibship size.

Consistent with other studies, we find that firstborn siblings have better educational outcomes than their later-born siblings. Figure 1 shows the magnitude of these associations. The top panel shows lower test scores for each successive birth order for all family sizes from 2-5 siblings. Most of these differences are present in the first test scores we observe (fifth grade), but the gaps do grow modestly from the last scores in our data (ninth grade) (see Supplementary Figure S2). The bottom panel shows similar patterns for educational attainment in the adult population. Figure 1 also shows that birth order differences increase when adjusted for maternal age.

**Fig. 1.**
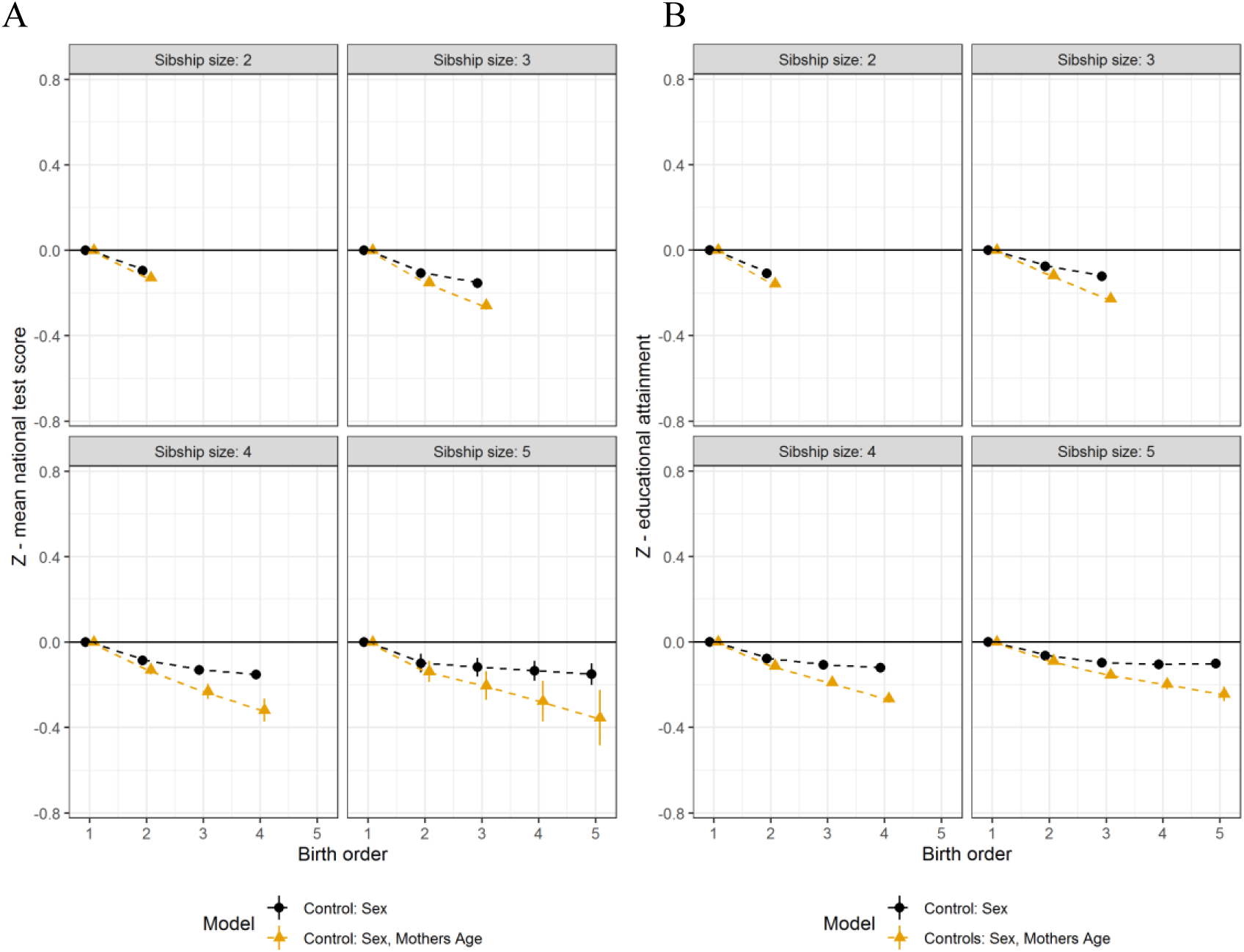
Birth order and educational differences in the population. **A**,**B** Results from family-fixed effects linear regression models run separately by sibship size, with controls for sex and maternal age and cluster-robust standard errors. Firstborns serve as the reference category. All point estimates presented with 95 % CI. In **A)** children part of the population (N=301,795) where the outcome is the mean of national test score standardized within test, year of test, and birth cohort. **B)** Parental part of the population (N=2,067,878), where the outcome is educational attainment at age 30, standardized within birth cohort.

### Polygenic score

Figure 2 shows the relationship between birth order and the polygenic score for educational attainment among families with 2 and 3 children, which are the vast majority of sibships in Norway. Sample sizes for this and subsequent analyses using genetic data are much smaller than those used in the previous figure as these data are only available for a portion of the population (see Methods). We provide separate results for the child sample (panels a and b; N = 24,507; 2,705) and their parents (panels c and d, N = 43,316; 3,897). Within each sample, we provide estimates for models comparing siblings both between families and within families. In addition to sex and cohort, between-family models adjust for the ten principal components of the GWAS data to address potential confounding by ancestral differences (i.e., population stratification).

**Figure 2.**
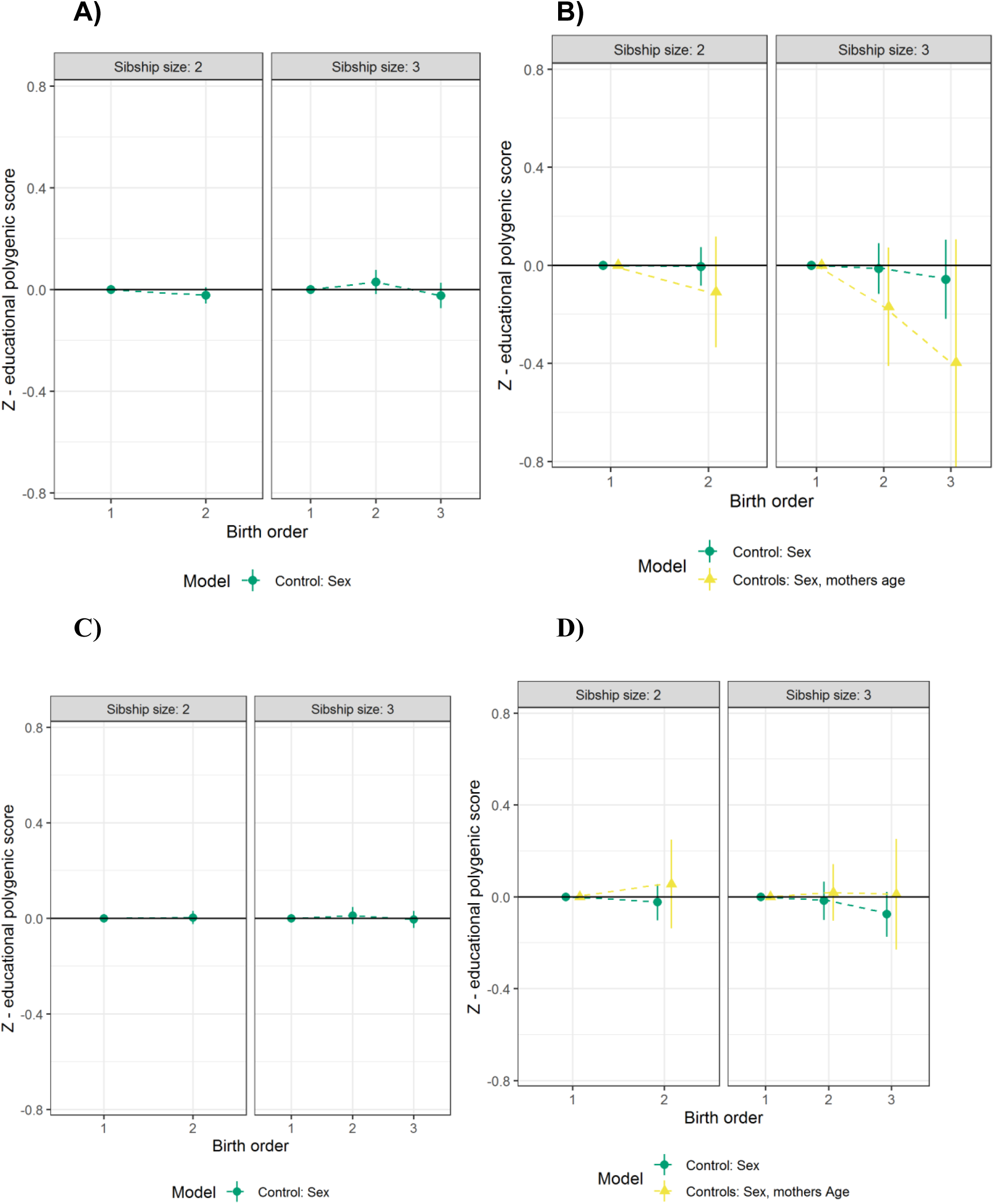
Polygenic score and birth order. **A, B, C, D**, Association between educational attainment polygenic score and birth order. Results from linear regression models run separately by sibship size, with controls for sex. Firstborns serve as the reference category. All point estimates presented with 95 % CI. In **A**,**C**, between-family estimate, adjusted for 10 principal components. In **B**,**D**, family-level fixed effects models with cluster-robust standard errors. In **A**,**B**, children part of the sample (N=24,507 (a); 2,705 (b));. **C**,**D**, parental part of the sample (N = 43,316 (c); 3,897 (d)).

In all panels in Figure 2, confidence intervals overlap zero for birth orders two and three, meaning that we find no differences in the polygenic score for educational attainment among different birth orders. Consequently, the genetic differences captured by that polygenic score cannot explain the observed relationship between birth order and educational outcomes. We also looked specifically at whether a child’s polygenic score for educational attainment predicted whether parents had another child, and we did not find any difference (Supplementary Table S1).

### Birth weight and birth length

In Figure 3, we turn to birth order differences in birth weight and birth length, available in the child part of the sample only (N=2,705). We adjust for sex, gestational age, and the polygenic score for education.

**Fig. 3.**
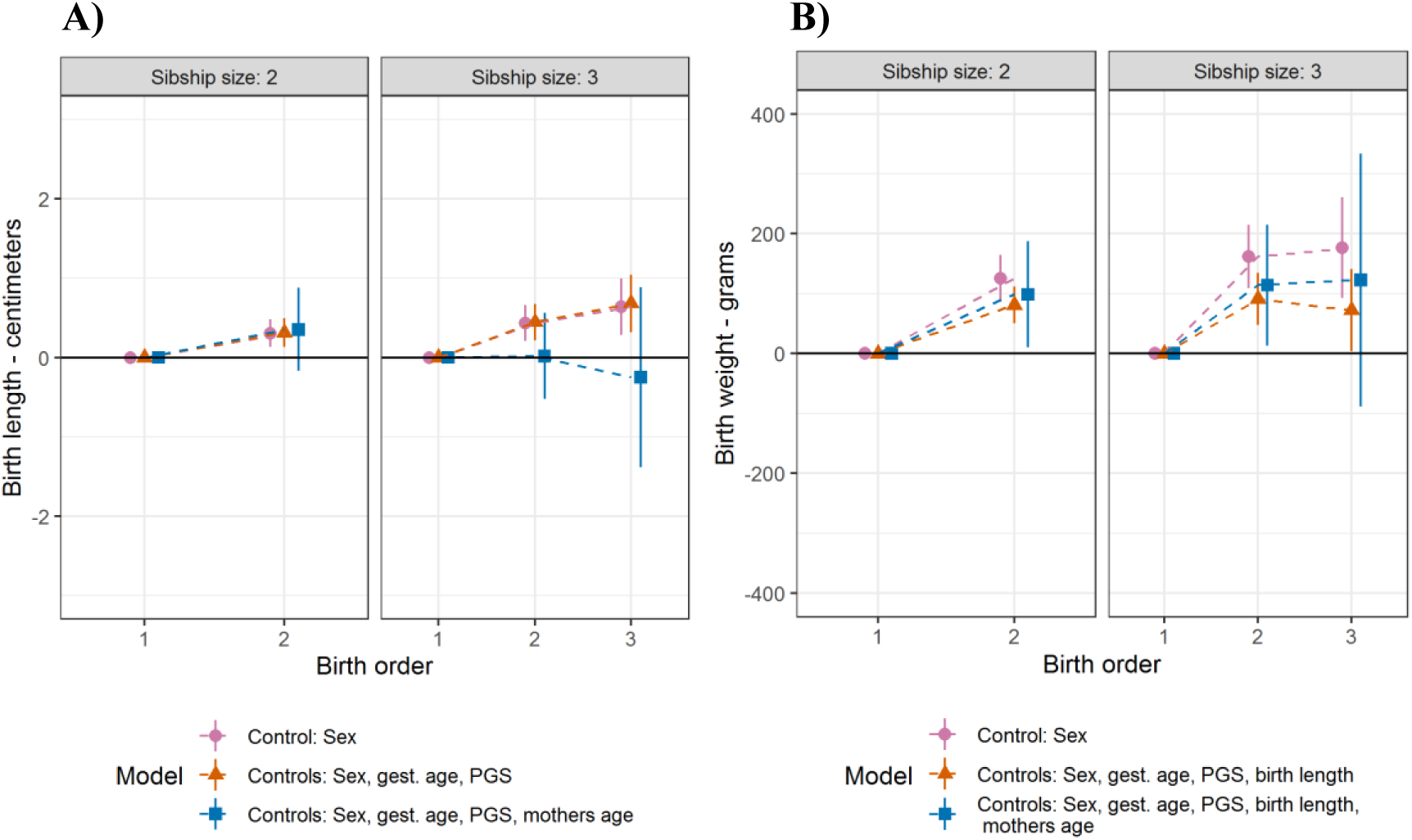
Birth length, birth weight and birth order. **A, B** Results from family-level fixed effects linear regressions run separately by sibship size, with dummies for birth order. Children part of the sample (N=2,705). Cluster robust standard errors, 95 % CI. Firstborns serve as the reference category. In **A**, birth length, with different control variables, pink point: sex; orange triangle: sex, gestational age, educational attainment polygenic score; blue rectangle: sex, gestational age, educational attainment polygenic score, mothers age at birth. In **B**, birth weight, with controls, pink point: sex; orange triangle: sex, gestational age, educational attainment polygenic score and birth length; blue rectangle: sex, gestational age, educational attainment polygenic score, birth length, mothers age at birth.

On average, later-born children are both longer and heavier than firstborn children. Differences in length account for roughly half of the difference in weight. Further analyses indicate that the differences in mean length or weight are not due to differences only in especially low-or high-weight births (see Supplementary Figure A3). We find nearly identical results in the between-family models (see Supplementary Figure S4). If higher birth weight indicates an advantaged pre-natal environment, then it appears that later born children are actually advantaged in this regard relative to their first born siblings, despite their lower ultimate achievement.

### Maternal age and birth spacing

We considered also the possibility that birth order differences were due to maternal age, as maternal age has been variously posited to influence cognitive development via both biological and social mechanisms. To confound an observed firstborn advantage, mothers age would need to be inversely associated with test scores. When we simultaneously model within-and between-family effects, we find maternal age to be positively associated with test scores between families and even more so within families (see Supplementary Table S2). Consequently, as noted, the birth order differences presented in Figure 1 actually increase after accounting for maternal age. Within families, sibling differences in maternal age are equivalent to the birth spacing intervals. Our finding is thus that birth order differences get smaller as both maternal age and the spacing between births increases. While this pattern is consistent with explanations of birth order differences rooted in differential parental investment and overall intellectual climate in the family, it is not consistent with ideas of there being in utero, mutational, or other biological advantages to being born to a younger mother.

### Educational achievement and attainment with controls

In Figure 4, we show birth order differences within families after adjusting for all the aforementioned measures simultaneously: sex, gestational age, birth weight, birth length, and mothers age at birth, as well the polygenic score for education. We show results for birth order differences in educational achievement (children, panel a, N=2,933) and educational attainment at age 30 (parents, panel b, N = 3,365). In all models, point estimates are negative for later-born children compared to firstborn. Although controlling for mothers’ age at birth increases the uncertainty of estimates and confidence intervals overlap zero (blue rectangle in Fig. 4), all point estimates remain negative, and some again increase.

**Fig. 4.**
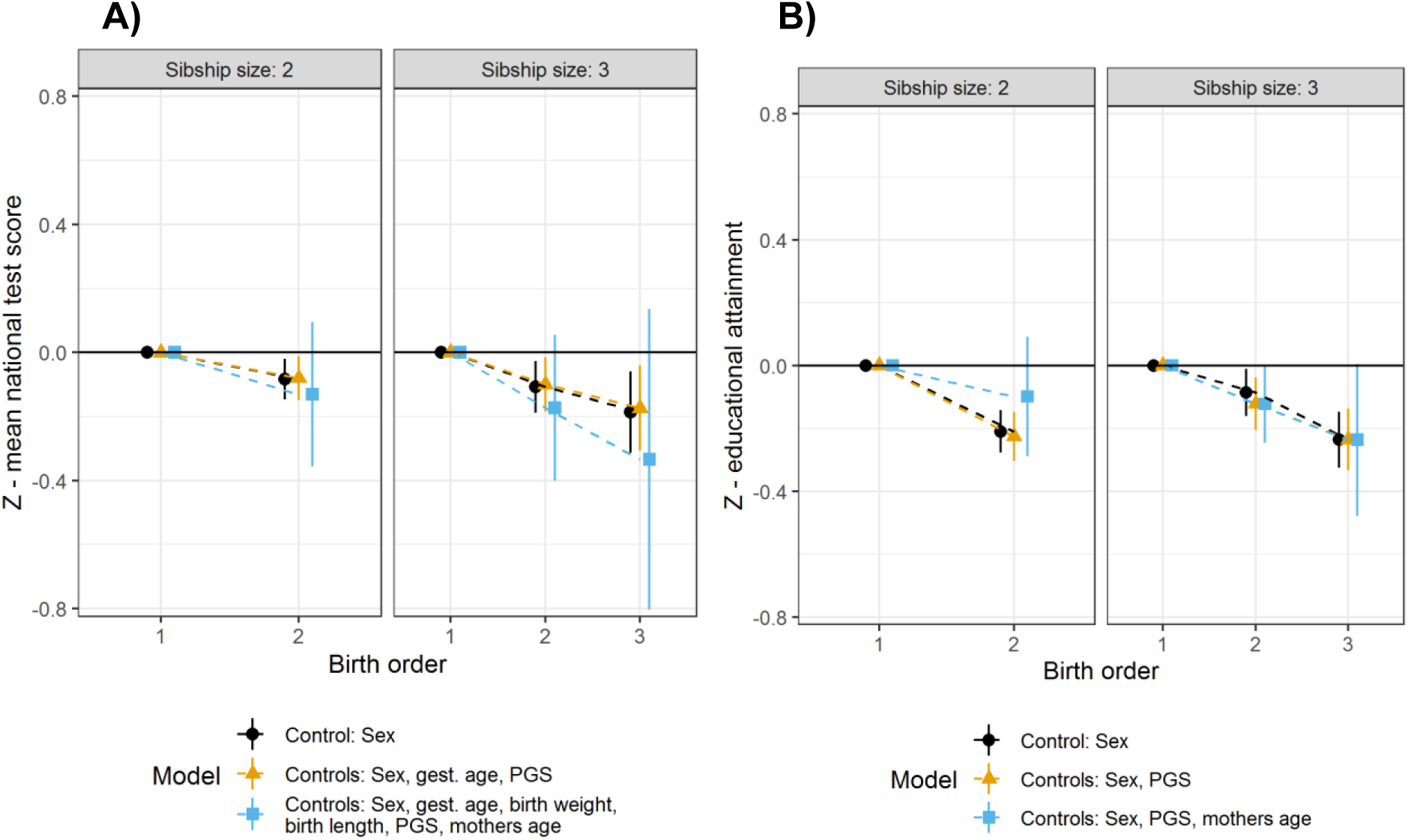
Educational achievement/attainment and birth order. **A**,**B** Results from family-level fixed effects linear regressions run separately by sibship size, with dummies for birth order where firstborns are the reference. Cluster robust standard errors, 95 % CI. In A, children part of the sample (N = 2,933) with mean of national test score as the outcome. Controls for sex (black circle); sex, gestational age, educational attainment polygenic score (yellow triangle); sex, gestational age, birth weight, birth length, educational attainment polygenic score (blue rectangle). In B, parental part of the sample (N = 3,365) with educational attainment at age 30 as the outcome. Controls for sex (black circle); sex, educational attainment polygenic score (yellow triangle), and sex, educational attainment polygenic score, mothers age (blue rectangle).

Taken together, these analyses rule out various scenarios by which birth order differences in educational outcomes are either confounded or the result of genetic or pre-natal causes. The availability of multigenerational genotyped data with extensive phenotyping does not contradict an understanding of birth order differences as originating in post-natal environments.

### Interrogating Environmental Origins

Given this, we next turned to see what additional clues the genomic data may offer for how these environmental influences operate. First, we examined whether birth order differences varied by polygenic score. Muslimova et al (45) posit that birth order differences may be strongest among children with the highest genetic score for achievement, as these children may be able to capitalize best on increased parental investment. In the child sample, we observed no difference (see Supplementary Table S3 and Supplementary Figure S5). In the adult sample, we find a borderline statistically significant pattern opposite the expected direction: a smaller firstborn advantage among participants with higher polygenic scores. If that is true, higher genetic potential could mitigate the processes that produce birth order differences. Either way, there is no indication in our data that firstborn advantage only or more strongly exists among those with higher polygenic scores.

Second, we also considered whether, net of polygenic score, birth order differences are moderated by family background. Contradictory hypotheses have been proposed to this end. One is that advantaged environments would have greater disparities in effective parental investment, implying larger birth order differences in those families. The other is that advantaged environments would have more compensatory investment in lower-performing children, yielding smaller birth order differences (see 46). Our results here were consistent with the former of these scenarios (Supplementary Table S4): differences were larger in families in which parents had higher educational attainment, including within-families and net of polygenic score. But, these differences by family background were small compared to the overall birth order difference (Supplementary Figure S6)

Third, we examined the influence of other family member’s polygenic scores on child achievement, net of the child’s own score. Recent findings of “genetic nurture” have documented relationships between non-transmitted parental alleles and child attainment and achievement (47–49). Given that mothers are typically more involved in childrearing than fathers--not infrequently to a substantial degree--the finding of some past research that maternal non-transmitted alleles matter more for attainments than paternal non-transmitted alleles would not be surprising (50). We did not find this pattern in our data (see Supplementary Table S5). Also, when we include the polygenic score for the siblings, the sibling score accounts for more than half of the magnitude of the difference in achievement that had been attributed to parental polygenic scores. Net of one another, the relationship between sibling polygenic score and achievement is nearly a quarter as large as that of a child’s own score (Supplementary Table S5). These results point to the importance of considering that genetic nurture may reflect sibling influence to a greater extent than has been previously appreciated.

## Discussion

Our starting point for this investigation was the common presumption that birth order differences in educational outcomes reflect social environmental rather than genetic or prenatal mechanisms. Specifically regarding genetic differences, there are multiple potential reasons for genetic variation by birth order. Historically the problem has been exacerbated by many studies using between-family samples and methods (51), but the possibility of genetic differences remains even in studies that compare siblings in the same family. The only way to assess decisively whether genetic differences may confound birth order studies is with data that contains genetic and comprehensive phenotypic information on many siblings. Using data that spans two generations of Norwegian siblings, we show no genetic differences by birth order as captured by polygenic scores for educational attainment. Similar preliminary findings have recently been reported using data from the United Kingdom (45).

We also examined indicators of possible *in utero* origins of birth order differences: birth weight, birth length, and maternal age. Our results suggest that none of these provide any leverage for explaining the robust advantage that earlier-born children have in test scores and ultimate educational attainment in our data. Indeed, later-born children are advantaged in birth weight and birth length, consistent with some earlier research (29, 52, 53), and so if anything they might have better in utero environments, and yet have worse educational outcomes.

These results strengthen the conclusion that birth order differences in educational outcomes mainly originate in post-birth environments (6, 39). Another implication of our findings is that various methodologies that use siblings in causal inferences, i.e. studies using polygenic scores within-family to alleviate population stratification (54, 55), and social science studies using siblings to capture the omnibus effect of (social background), are not confounded in ways they would be if birth order differences were due to genetic differences.

Given that birth order differences seem not related to genetic factors, nor do they appear to be due to in utero environments, they constitute one of the clearest-cut examples of postnatally-induced social inequality within families. Understanding how they come about offers a window that may be more generally instructive for mechanisms that lead to inequalities among people with similar familial and sociodemographic backgrounds. To this end, we find that the birth order differences in test scores are already mostly present at our earliest time of observation (5th grade, age 9/10) and grow only modestly from then to 9th grade. This clarifies that the environmental differences associated with birth order differences largely manifest in the first decade of life. The difference in educational attainment among adults is similar in magnitude, suggesting that these differences resulting from childhood environments have lifelong effects.

The two main environmental theories on birth order effects are linked to parental resources and sibling interactions. According to the resource dilution theory (12, 56, 57), economic and parental resources deplete as more household members arrive. In societies where resources are abundant like Norway, cultural resources are thought to be more important than economic (Conley 2005). According to the theory, as earlier-born receive more of the parental cultural resources such as personal attention and help with homework than later-borns do, they excel in educational performance. While this pattern is consistent with the finding that birth order differences decline with increased spacing between births, it is not consistent with our results that maternal genetic endowments do not matter anymore than paternal endowments, given that mothers usually contribute more to child-rearing than fathers. Other studies have found stronger effects of maternal genetics (50), so more research will be needed to reach any decisive conclusion.

As for sibling interactions, these have been most prominently raised in variations of “confluence theory”, which proposes that siblings generally have negative effects on one another’s cognitive development (13, 14, 58). Because firstborns have less exposure to being reared with siblings, and may also benefit from the opportunity to teach younger siblings, the negative influence of siblings is proposed to be least for firstborns. Our finding that the apparent influence of parental alleles shrinks markedly when sibling polygenic scores are included, and that sibling polygenic scores are significant associated with child outcomes net of a child’s own score, indicates that some of what has hitherto been called “genetic nurture” could in fact be “sibling genetic nurture.” Recent enthusiasm for using sibling-based GWAS to purge GWAS studies of indirect genetic effects assumes that these indirect genetic effects are primarily due to parents (59). Because the approach does not account for indirect effects via siblings, the possibility that sibling genetic effects on achievement may be larger than commonly supposed (60) is important to resolve for evaluating the success of sibling-based GWAS.

Sibling effects have received less attention as compared to parental resources when it comes to explaining birth order effects. That said, Gibbs et al (61) propose a conditional resource dilution (CRD) model, opening up for institutional- and family-level variation influencing birth order effects. In the CRD, brothers and sisters, and especially older siblings, may provide, rather than compete for resources. “Sibling genetic nurture” raises the possibility that these intersibling consequences for familial environments vary by sibling genotype, with genetic factors increasing achievement potential having positive spillovers for siblings.

Our study has some limitations. First, polygenic scores are based on common alleles, and we are not able to tell if there are differences in rarer variants. That said, If rarer variants in the form of de novo mutations were importantly associated with birth order, the most obvious explanation would have involved parental age, and this is contradicted by our findings regarding maternal age. Second, our proxies for in utero environments are partial, such that unmeasured consequential differences might still exist. Similarly, even though post-natal accounts of birth order differences focus on social environments, our design does rule out postnatal differences due to physiological causes or other, non-social phenomena. Third, the sample size for the within-family analyses is limited, as we have relatively few families with two or more full siblings. Statistical power curtails our present capacity to draw more decisive conclusions for some of our analyses, and we cannot rule out type II errors. Fourth, polygenic scores for educational attainment still only account for a limited portion of the overall heritability of educational attainment, and they will likely improve in the future. Last, we have no direct measures of parental and sibling behaviors, which obviously thwarts the ability to interrogate either as putative causes. Amidst the excitement for what incorporating genomic information may bring to the social sciences, we should not lose sight that the available phenotyping of population datasets often still leaves much to be desired.

In addition, even though birth order differences seem like a unary puzzle, the importance of different environmental mechanisms may shift in different societal and historical contexts. For the cohorts in our child sample, the vast majority of Norwegian children attend child care from age 1, with the overall attendance being 90 % for children aged 1-5 in 2010 (62). In such a context, mothers may be spending less time with their children than perhaps at any point in history, which might mean our data could be unpropitious for observing a maternal role in birth order effects. Scandinavian data has often been used in research on family dynamics research because of these countries’ population registers. Societies with exceptional data may also be exceptional in other respects. Moreover, Scandinavian population register data sources do not contain large-scale data on actual behaviors, resources, and practices within families, and there may simply be no way of getting to the bottom of birth order effects without such richer data. While our results show that birth order differences are largely environmental, the question of what specific environments factors are involved remains far from resolved.

## Materials and Methods

### Population data and variables from administrative registries

The data in this paper are from several sources. We begin with data from administrative registries covering the full population of Norway. The registries are of very high quality, and do not suffer from attrition, and have few registration errors. From the Central Population Registry we identify all family linkages and demographic variables, like birth cohort, sibship size, birth order, and mothers age at birth. We identify sibship size and birth order according to birth year within each mother. We also identify any sibships with multiple births, and exclude them as the assignment of birth order is less clear cut, and family dynamics may be different in multiple birth sibships. To better mirror our genomic sample we remove individuals born outside of Norway. Current genomic methods do not allow for us to include persons of non-European ancestry, and we restrict the data based on the population registry based on this related criteria, i.e. we only include Norwegian-born to Norwegian parents. For the adult part of the population, we restrict birth cohorts 1945-1988.

We link the Central Population Registry to the National Educational database (NuDB) (63), also covering the full population. For the children part of our analysis, we use standardized tests as the outcome. From NuDB we use data from national tests conducted in 5th, 8th, and 9th grade in reading, mathematics, and English. We standardize each test to a z-score within test, year of test assessment, and birth cohort. Thereafter, we calculate a mean of all available test scores for each child, which serves as our outcome variable for the children in the sample. For the adult part of our analysis, the outcome is educational attainment at age 30 in years following the International Standard Classification of Education (ISCED 2011) (63, 64). The variable is continuous as we transform it to being measured in years, using how many years it takes to complete the level of education attained following normal progression, according to ISCED. We standardize this variable too to compare effect sizes between outcomes. After restricting our sample and removing people with missing information etc., our population-based data has an N of 301,795 for the child birth cohorts, and 2,067,878 for the adult birth cohorts.

### The Norwegian Mother, Father and Child Cohort Study (MoBa)

The prepared population data are linked to The Norwegian Mother, Father and Child Cohort Study (MoBa) (65, 66), which we use for the analysis with genomic and in-utero variables. MoBa is a population-based pregnancy cohort study conducted by the Norwegian Institute of Public Health. Participants were recruited from all over Norway from 1999-2008, with the sample unit being pregnancy. The women consented to participation in 41% of the pregnancies. The cohort includes 114,500 children, 95,200 mothers and 75,200 fathers. The current study is based on version 12 of the quality-assured data files. The establishment of MoBa and initial data collection was based on a license from the Norwegian Data Protection Agency and approval from The Regional Committees for Medical and Health Research Ethics. The MoBa cohort is now based on regulations related to the Norwegian Health Registry Act. The current study was approved by The Regional Committees for Medical and Health Research Ethics. Some 98 110 individuals in around 32 000 trios are currently genotyped. This interim release is known as “MoBa Genetics”, and is comprised of several separately genotyped and imputed batches due to partial funding for genotyping on a per-project basis. The current release is a merger of all subprojects after quality control and imputation. Details are available here: https://github.com/folkehelseinstituttet/mobagen.

MoBa-participants are sampled on pregnancy independent of previous pregnancies (67), meaning that children in MoBa could have siblings younger or older not included in the sample. However, as we identify family linkages and birth order from the population registry, we observe full sibship sizes and birth orders independently of the information in MoBa. The sampling of births within birth order is therefore random. As we construct sibship size variables from registries with observations from the birth of the parents and up until 2018, we are confident that we observe completed fertility histories for the vast majority of the sample, as most have reached ages where fertility rates are very low. For the parental part of the MoBa-sample, we use the central population registry to link them to their parents (i.e. the grandparents of children in MoBa). By doing this, we can establish sibship size, birth order, and other demographic information from the Central Population Registry, and use that to investigate birth order differences taking into account genomic information also for the parental part of the MoBa-sample. Our observational window from the NuDB register is up until 2018. In a few cases the parents in MoBa have yet to have reached age 30 at our latest observation year 2018. Here, we take the latest observed age available, the lowest age being 27.

### Variables from MoBa

MoBa contains genomic information for parents and children which allows us to create Polygenic Scores (PGS) for each individual based on a Genome-Wide-Association study (GWAS) for educational attainment of 1,1 million people (68). For the MoBa-sample we conducted quality control using PLINK (v 1.90). Before we perform quality control in PLINK, we remove families with any individuals born in countries outside of Europe, The United States, Canada, New Zealand, or Australia. We also remove families with any multipartnered fertility parents in MoBa. Thresholds for genotyping call rate were set to 98 %, minor allele frequency (MAF) 5 %, and deviations from Hardy-Weinberg equilibrium was <10^−4^ with mid p-value adjustment. We remove individuals with poor genotype quality with a threshold missing rate of 5 %, as well as those with heterozygosity rates which deviate ±3 standard deviations from the sample mean. We remove ancestry outliers in plink with an identity-by-state binominal test (PPC) set to 0.05, MAF 0.01 and compare the 1-5 nearest neighbors in the data. We remove families with a z-score for any neighbor below 4 standard deviations. We use KING software (69) to confirm that siblings are genetic siblings based on identity-by descent.

We use PRSice (70) to make polygenic scores for parents and children. We use all available SNPs, genome-wide significant or not, and clump with a 250 kb window. Clump-r2 was set to 0.1, excluding SNPs with higher linkage disequilibrium. The polygenic score is standardized.

In addition to genomic information, MoBa has several survey waves. Birth length and birth weight were self-reported from the mother when the child was 6 months old. Questionnaires and instrument documentation for the MoBa is available here: https://www.fhi.no/en/studies/moba/for-forskere-artikler/questionnaires-from-moba/

After doing quality control on the genomic data, and removing people according to what we have just described, 79 057 individuals are left in the sample, 29 815 children, 24 032 fathers and 25 210 mothers. However, while we use this sample for between-family analysis in some figures, we rely mostly on within-family analysis, i.e. family fixed-effects with clustering on mothers and robust standard errors. Here, our sample size is smaller, as at least two children born to the same mother are needed to estimate the models, and we need both siblings to be genotyped (N=2,933).

## Data availability

The consent given by the participants does not open for storage of data on an individual level in repositories or journals. Researchers who want access to data sets for replication should submit an application to. Access to data sets requires approval from The Regional Committee for Medical Research Ethics in Norway and a formal contract with MoBa. Code is available at https://osf.io/7j5ez/

## Acknowledgments

This research is part of the OPENFLUX project that has received funding from the European Research Council (ERC) under European Union’s Horizon 2020 research and innovation programme (ERC Consolidator Grant agreement No. 818420). The Norwegian Mother, Father and Child Cohort Study is supported by the Norwegian Ministry of Health and Care Services and the Ministry of Education and Research. We are grateful to all the participating families in Norway who take part in this on-going cohort study. We thank the Norwegian Institute of Public Health (NIPH) for generating high-quality genomic data. This research is part of the HARVEST collaboration, supported by the Research Council of Norway (#229624). We also thank the NORMENT Centre for providing genotype data, funded by the Research Council of Norway (#223273), South East Norway Health Authority and KG Jebsen Stiftelsen. We thank Per Minor Magnus, Pål Njølstad and Ole Andreassen who headed the forementioned projects. We further thank the Center for Diabetes Research, the University of Bergen for providing genotype data and performing quality control and imputation of the data funded by the ERC AdG project SELECTionPREDISPOSED, Stiftelsen Kristian Gerhard Jebsen, Trond Mohn Foundation, the Research Council of Norway, the Novo Nordisk Foundation, the University of Bergen, and the Western Norway health Authorities (Helse Vest). Isungset thank Robbee Wedow for making genomics less hard.

## Notes

### Competing Interest Statement

The authors have declared no competing interest.

## References

1. D. Conley, The Pecking Order (Vintage Books, 2005).

2. E. Turkheimer, Weak Genetic Explanation 20 Years Later. Perspect. Psychol. Sci. 11, 24–28 (2016).

3. R. Plomin, D. Daniels, Why are children in the same family so different from one another? Behav. Brain Sci. 10, 1–16 (1987).

4. J. M. Rohrer, B. Egloff, S. C. Schmukle, Examining the effects of birth order on personality. Proc. Natl. Acad. Sci. 112, 14224–14229 (2015).

5. L. Belmont, F. A. Marolla, Birth Order, Family Size, and Intelligence: A study of a total population of 19-year-old men born in the Netherlands is presented. Science (80-.). 182, 1096–1101 (1973).

6. P. Kristensen, T. Bjerkedal, Explaining the Relation Between Birth Order and Intelligence. Science (80-.). 316, 1717–1717 (2007).

7. S. E. Black, P. J. Devereux, K. G. Salvanes, The More the Merrier? The Effect of Family Size and Birth Order on Children’s Education*. Q. J. Econ. 120, 669–700 (2005).

8. L. Esposito, S. M. Kumar, A. Villaseñor, The importance of being earliest: birth order and educational outcomes along the socioeconomic ladder in Mexico. J. Popul. Econ. 33, 1069–1099 (2020).

9. J. R. Behrman, P. Taubman, Birth Order, Schooling, and Earnings. J. Labor Econ. 4, 121–150 (1986).

10. M. Bertoni, G. Brunello, Later-borns Don’t Give Up: The Temporary Effects of Birth Order on European Earnings. Demography 53, 449–470 (2016).

11. J.-Y. K. Lehmann, A. Nuevo-Chiquero, M. Vidal-Fernandez, The Early Origins of Birth Order Differences in Children’s Outcomes and Parental Behavior. J. Hum. Resour. 53, 123–156 (2018).

12. J. Blake, Family Size and Achievement (University of California Press, 1989).

13. R. Zajonc, G. Markus, Birth Order and Intellectual Development. Psychol. Rev. 82, 74–88 (1975).

14. R. B. Zajonc, Validating the Confluence Model. Psychol. Bull. 93, 457–480 (1983).

15. K. Hyeokmoon, D. O. Martschenko, K. P. Harden, T. A. Diprete, P. D. Koellinger, Genetic Fortune : Winning or Losing Education, Income, and Health (2020).

16. K. P. Harden, The Genetic Lottery - Why DNA Matters for Social Equality (Princeton University Press, 2021).

17. P. Engzell, F. C. Tropf, Heritability of education rises with intergenerational mobility. Proc. Natl. Acad. Sci. U. S. A. 116, 25386–25388 (2019).

18. A. Björklund, K. G. Salvanes, Education and Family Background. Mechanisms and Policies. IZA Discuss. Pap. (2010) https://doi.org/10.1016/B978-0-444-53429-3.00003-X.

19. E. Tucker-Drob, M. Rhemtulla, K. Harden, E. Turkheimer, D. Fask, Emergence of a Gene-by-Socioeconomic Status Interaction on Infant Mental Ability from 10 Months to 2 Years. Psychol Sci 22, 125–133 (2011).

20. P. R. Jansen, et al., Polygenic scores for schizophrenia and educational attainment are associated with behavioural problems in early childhood in the general population. J. Child Psychol. Psychiatry Allied Discip. 59, 39–47 (2018).

21. J. Wertz, et al., Using DNA From Mothers and Children to Study Parental Investment in Children’s Educational Attainment. Child Dev. 00, cdev.13329 (2019).

22. A. Breinholt, D. Conley, Child-Driven Parenting: Differential Early Childhood Investment by Offspring Genotype. NBER Work. Pap. (2020).

23. R. Linnakaari, et al., Trends in the incidence, rate and treatment of miscarriage - Nationwide register-study in Finland, 1998-2016. Hum. Reprod. 34, 2120–2128 (2019).

24. J. S. Cohain, R. E. Buxbaum, D. Mankuta, Spontaneous first trimester miscarriage rates per woman among parous women with 1 or more pregnancies of 24 weeks or more. BMC Pregnancy Childbirth 17, 1–7 (2017).

25. A. García-Enguídanos, M. E. Calle, J. Valero, S. Luna, V. Domínguez-Rojas, Risk factors in miscarriage: A review. Eur. J. Obstet. Gynecol. Reprod. Biol. 102, 111–119 (2002).

26. D. Malaspina, et al., Paternal age and intelligence: Implications for age-related genomic changes in male germ cells. Psychiatr. Genet. 15, 117–125 (2005).

27. D. Almond, J. Currie, Killing Me Softly: The Fetal Origins Hypothesis. J. Econ. Perspect. 25, 153–172 (2011).

28. J. W. Foster, S. J. Archer, Birth Order and Intelligence: an Immunological Interpretation. Percept. Mot. Skills, 79–93 (1979).

29. A. A. Brenøe, R. Molitor, Birth order and health of newborns: What can we learn from Danish registry data? J. Popul. Econ. 31, 363–395 (2017).

30. S. E. Black, P. J. Devereux, K. G. Salvanes, FROM THE CRADLE TO THE LABOR MARKET? THE EFFECT OF BIRTH WEIGHT ON ADULT OUTCOMES*. Q. J. Econ. 122 (2007).

31. D. Conley, N. G. Bennett, Is Biology Destiny ? Birth Weight and Life Chances. Am. Sociol. Rev. 65, 458–467 (2000).

32. B. C. Young, R. J. Levine, S. A. Karumanchi, Pathogenesis of preeclampsia. Annu. Rev. Pathol. Mech. Dis. 5, 173–192 (2010).

33. S. B. Gumusoglu, A. S. S. Chilukuri, D. A. Santillan, M. K. Santillan, H. E. Stevens, Neurodevelopmental Outcomes of Prenatal Preeclampsia Exposure. Trends Neurosci. 43, 253–268 (2020).

34. C. S. Wu, et al., Health of children born to mothers who had preeclampsia: a population-based cohort study. Am. J. Obstet. Gynecol. 201, 269.e1-269.e10 (2009).

35. A. Kong, et al., Rate of de novo mutations and the importance of father’s age to disease risk. Nature 488, 471–475 (2012).

36. J. L. Taylor, et al., Paternal-age-related de novo mutations and risk for five disorders. Nat. Commun. 10, 1–9 (2019).

37. J. M. Goldmann, J. A. Veltman, C. Gilissen, De Novo Mutations Reflect Development and Aging of the Human Germline. Trends Genet. 35, 828–839 (2019).

38. G. A. Sartorius, E. Nieschlag, Paternal age and reproduction. Hum. Reprod. Update 16, 65–79 (2009).

39. K. Barclay, Birth order and educational attainment: Evidence from fully adopted sibling groups. Intelligence 48, 109–122 (2015).

40. M. A. Bellis, K. Hughes, S. Hughes, J. R. Ashton, Measuring paternal discrepancy and its public health consequences. J. Epidemiol. Community Health 59, 749–754 (2005).

41. M. Lillehagen, M. A. Isungset, New Partner, New Order? Multipartnered Fertility and Birth Order Effects on Educational Achievement. Demography 57, 1625–1646 (2020).

42. E. Thomson, T. Lappegård, M. Carlson, A. Evans, E. Gray, Childbearing Across Partnerships in Australia, the United States, Norway, and Sweden. Demography 51, 485–508 (2014).

43. N. M. Davies, et al., Within family Mendelian randomization studies. Hum. Mol. Genet. 28, R170–R179 (2019).

44. Ø. Kravdal, Taking birth year into account when analysing effects of maternal age on child health and other outcomes: The value of a multilevelmultiprocess model compared to a sibling model. Demogr. Res. 40, 1249–1290 (2019).

45. D. Muslimova, H. Van Kippersluis, C. A. Rietveld, S. Von Hinke, S. F. W. Meddens, Dynamic complementarity in skill production : Evidence from genetic endowments and birth order. Tinbergen Inst. Discuss. Pap., 1–46 (2020).

46. J. Price, Parent-Child Quality Time: Does Birth Order Matter? J. Hum. Resour. 43, 240–265 (2008).

47. M. A. Isungset, et al., Social and Genetic Effects on Educational Performance in Early Adolescence. NBER Work. Pap., 1–27 (2021).

48. T. C. Bates, et al., The Nature of Nurture: Using a Virtual-Parent Design to Test Parenting Effects on Children’s Educational Attainment in Genotyped Families. Twin Res. Hum. Genet. 21, 73–83 (2018).

49. B. Wang, et al., Robust genetic nurture effects on education: A systematic review and meta-analysis based on 38,654 families across 8 cohorts. Am. J. Hum. Genet. 108, 1780–1791 (2021).

50. A. Kong, et al., The nature of nurture: Effects of parental genotypes. Science (80-.). 359, 424–428 (2018).

51. J. L. Rodgers, What Causes Birth Order-Intelligence Patterns? The Admixture Hypothesis, Revived. Am. Psychol. 56, 505–510 (2001).

52. P. Magnus, K. Berg, T. Bjérkedal, The association of parity and birth weight: testing the sensitization hypothesis. Early Hum. Dev. 12, 49–54 (1985).

53. G. K. Swamy, S. Edwards, A. Gelfand, S. A. James, M. L. Miranda, Maternal age, birth order, and race: Differential effects on birthweight. J. Epidemiol. Community Health 66, 136–142 (2012).

54. T. T. Morris, N. M. Davies, G. Hemani, G. D. Smith, Population phenomena inflate genetic associations of complex social traits. Sci. Adv. 6 (2020).

55. K. P. Harden, P. D. Koellinger, Using genetics for social science. Nat. Hum. Behav. (2020) https://doi.org/10.1038/s41562-020-0862-5.

56. J. Blake, Family Size and the Quality of Children. Demography 18, 421–442 (1981).

57. J. Blake, Number of siblings, family background, and the process of educational attainment. Soc. Biol. 33, 5–21 (1986).

58. R. B. Zajonc, Family configuration and intelligence. Science (80-.). 192, 227–236 (1976).

59. L. J. Howe, et al., Within-sibship GWAS improve estimates of direct genetic effects. bioRxiv (2021) https://doi.org/10.1101/2021.03.05.433935.

60. A. I. Young, et al., Mendelian imputation of parental genotypes for genome-wide estimation of direct and indirect genetic effects. bioRxiv, 1–19 (2020).

61. B. G. Gibbs, J. Workman, D. B. Downey, The (Conditional) Resource Dilution Model: State- and Community-Level Modifications. Demography 53, 723–748 (2016).

62. R. H. Kitterød, E. H. Nymoen, J. Lyngstad, Endringer i bruk av barnetilsyn fra 2002 til 2010. Tabellrapport [Changes in use of child care from 2002 to 2010] (2012).

63. N. Barrabés, G. K. Østli, “Norwegian Standard Classification of Education 2016” (2017).

64. UNESCO Institute for Statistics, “The International Standard Classification of Education (ISCED)” (2012) https://doi.org/10.1007/BF02207511.

65. P. Magnus, et al., Cohort Profile Update: The Norwegian Mother and Child Cohort Study (MoBa). Int. J. Epidemiol. 45, 382–388 (2016).

66. L. Paltiel, et al., The Biobank in the Norwegian Mother and Child Cohort Study. Nor. Epidemiol. 16, 29–35 (2014).

67. K. S. Rønningen, et al., The biobank of the Norwegian mother and child cohort Study: A resource for the next 100 years. Eur. J. Epidemiol. 21, 619–625 (2006).

68. J. J. Lee, et al., Gene discovery and polygenic prediction from a genome-wide association study of educational attainment in 1.1 million individuals. Nat. Genet. 50, 1112–1121 (2018).

69. A. Manichaikul, et al., Robust relationship inference in genome-wide association studies. Bioinformatics 26, 2867–2873 (2010).

70. S. W. Choi, T. S.-H. Mak, P. F. O’Reilly, Tutorial: a guide to performing polygenic risk score analyses. Nat. Protoc. (2020) https://doi.org/10.1038/s41596-020-0353-1.

