## Supplementary material for "Birth Order Differences in Education Originate in Post-Natal Environments": SI

Martin Arstad Isungset (ORCID: 0000-0001-9316-3279)
Jeremy Freese
Ole A. Andreassen (ORCID: 0000-0002-4461-3568)
Torkild Hovde Lyngstad (ORCID: 0000-0001-7830-9305)

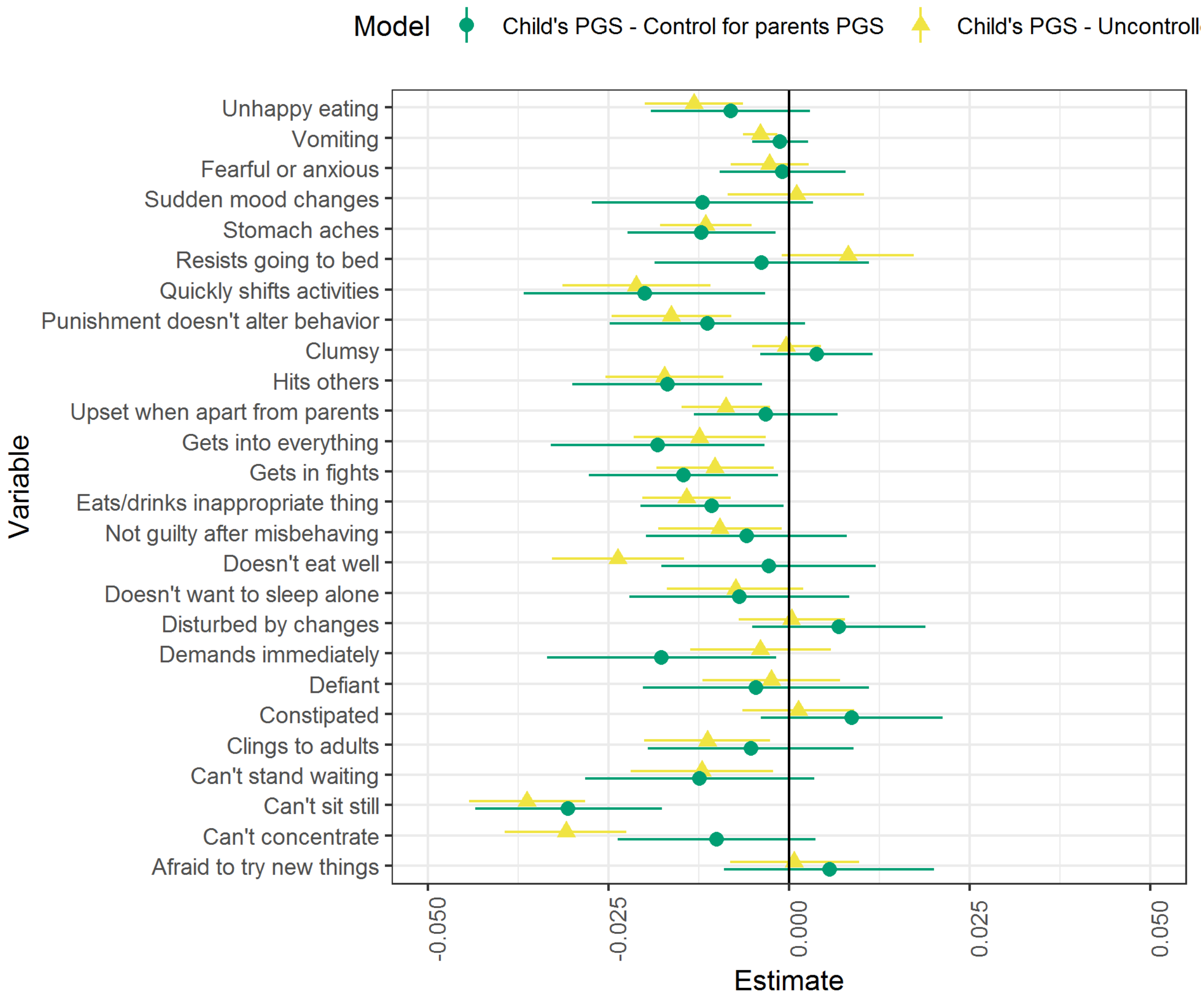


**Fig. S1.** Polygenic score and items from Child Behaviour Checklist (CBCL). Measured at 36 months. Full instrument documentation available here: https://www.fhi.no/globalassets/dokumenterfiler/studier/den-norske-mor-far-og-barn--undersokelsenmoba/instrumentdokumentasjon/instrument-documentation-q6.pdf

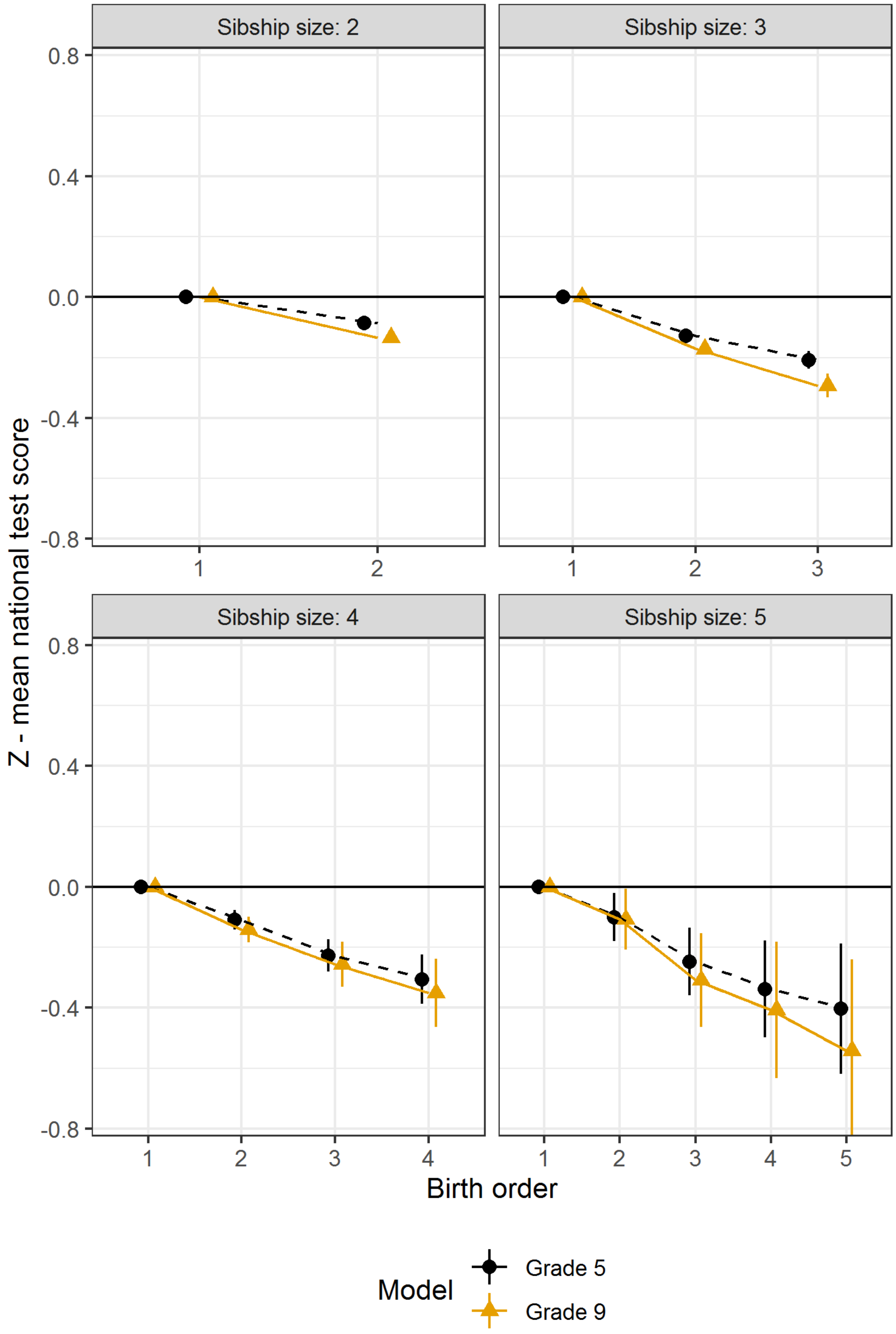


**Fig. S2.** Birth order and educational achievement in 5th and 9th grade - population. A,b Results from family-fixed effects linear regression models run separately by sibship size, with controls for sex and maternal age and cluster-robust standard errors. Firstborn serve as reference category. All point estimates presented with 95 % CI. N = 301,795

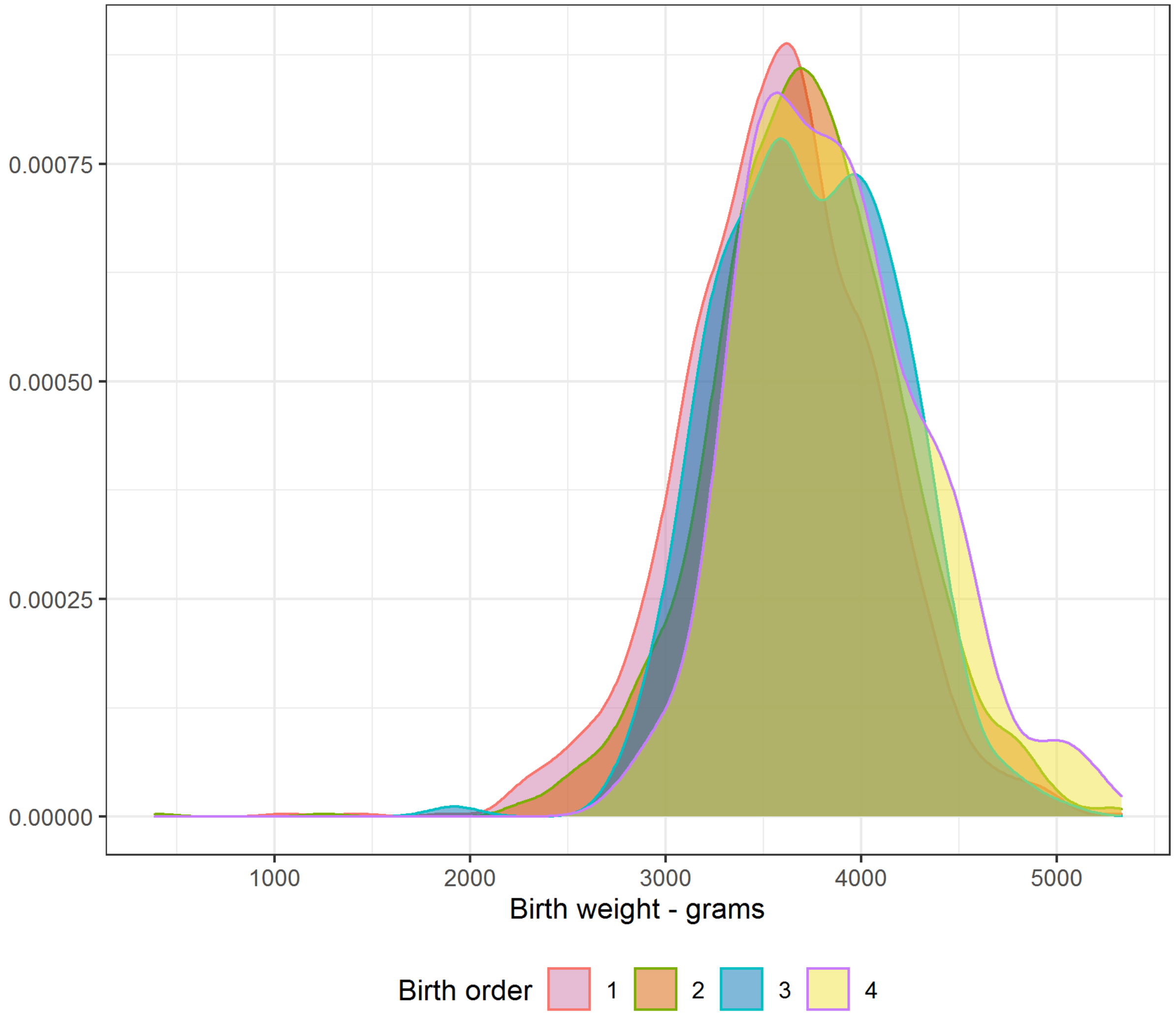


**Fig. S3.** Birth order birth weight - distribution plot

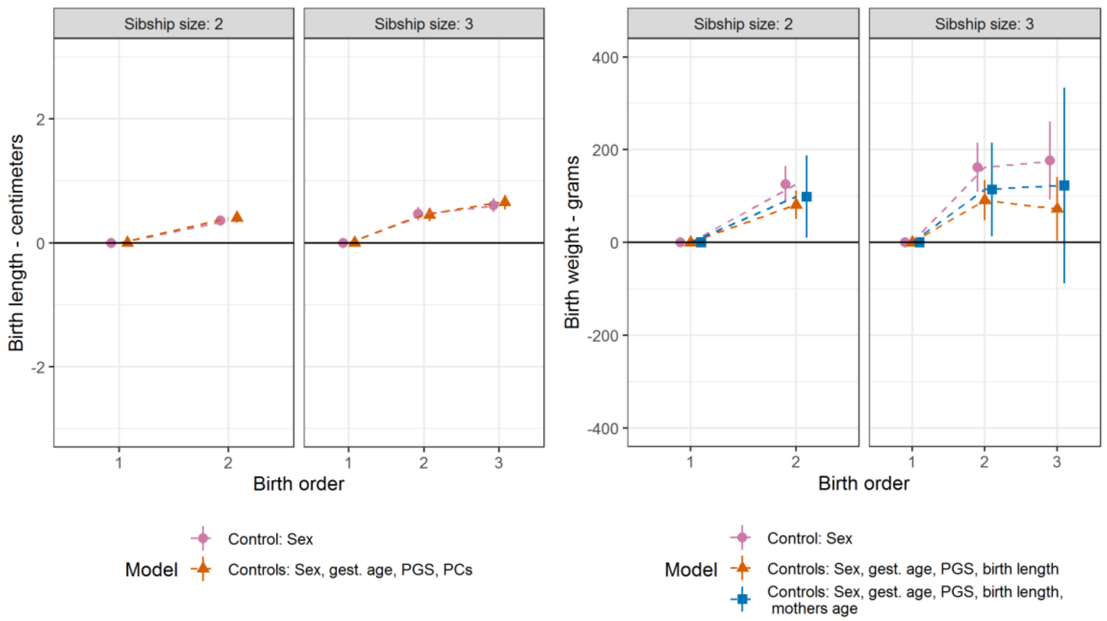


**Fig. S4.** Birth length, birth weight and birth order. A,b Results from linear regressions run separately by sibship size, with dummies for birth order. Children part of the sample (N = 8,188). Cluster robust standard errors, 95 % CI. Firstborns serve as reference category. In **a,** birth length, with different control variables, pink point: sex; orange triangle: sex, gestational age, educational attainment polygenic score, and principal components. In **b,** birth weight, with controls, pink point: sex; orange triangle: sex, gestational age, educational attainment polygenic score, and birth length

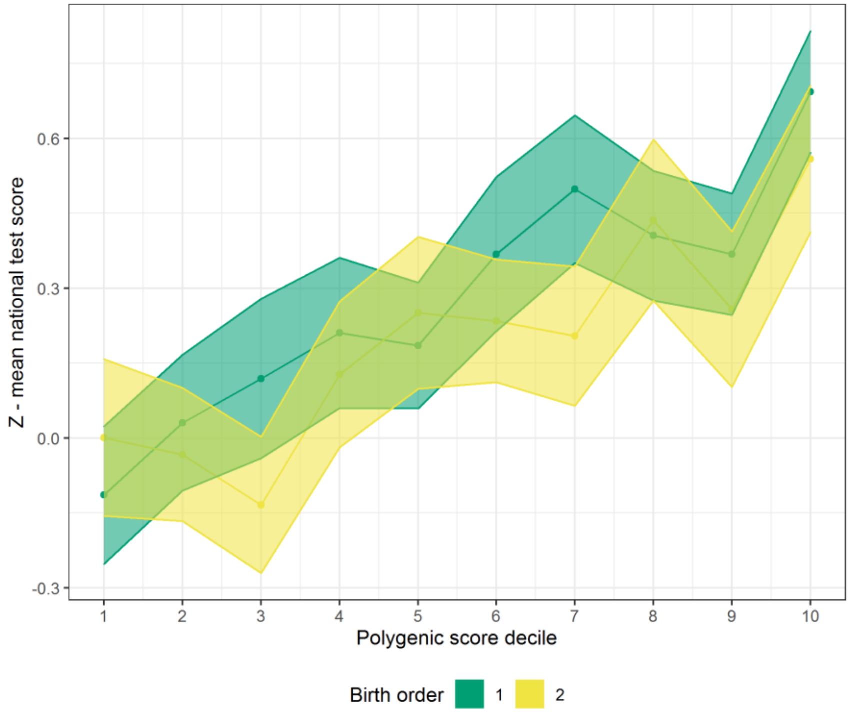


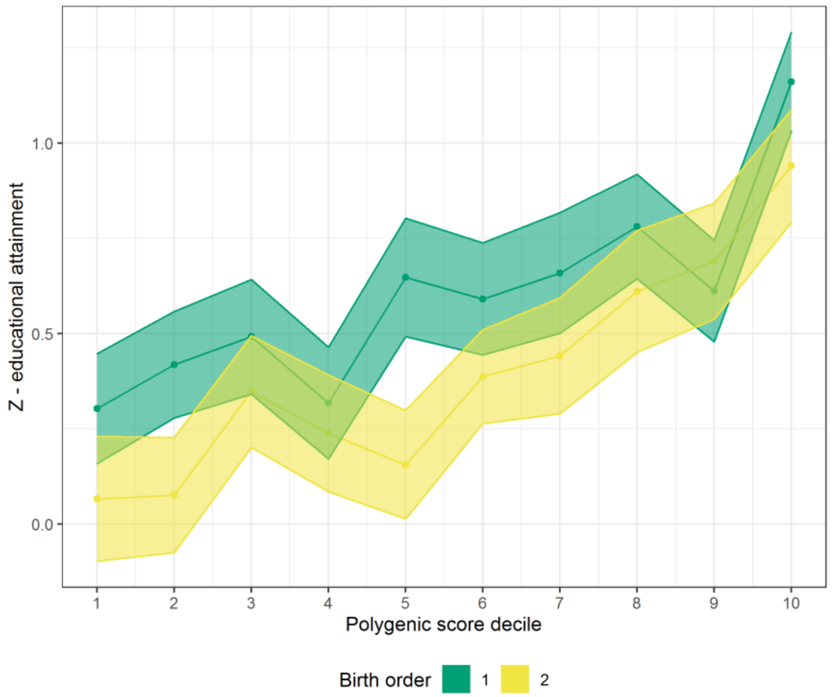


**Fig. S5.** Polygenic Score decile, birth order and educational achievement/attainment. Sibship size two only. We use 83.4% confidence intervals used for point estimates so that non-overlapping estimates indicates a difference between estimates at 95% confidence (1)

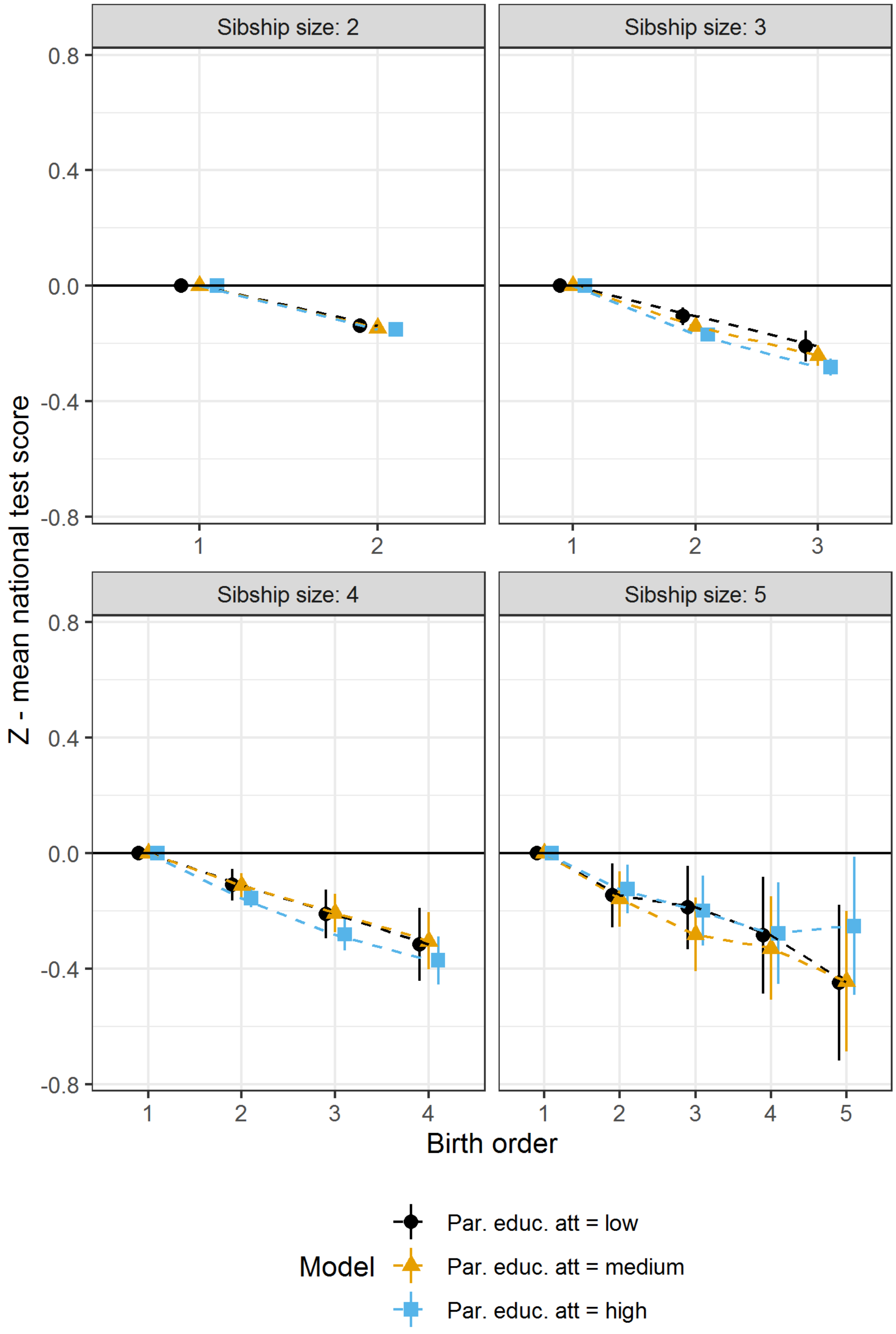


**Fig S6.** Parental educational attainment and birth order differences. Children - population. Parental educational attainment is a average parental educational attainment measured in years and then divided to treciles.

**Table S1.** Fertility decision x previous born child's PGS. Outcomes is having had another child or not

| | *Dependent variable:* | | |
|---|---|---|---|
| | Dummy - Had another child = 1 | | |
| | (1) | (2) | (3) |
| Child PGS Minus Avg. Parents PGS | 0.001 | | |
| | (-0.008, 0.009) | | |
| Child PGS Minus Mother PGS | | -0.005 | |
| | | (-0.012, 0.002) | |
| Child PGS Minus Father PGS | | | 0.005 |
| | | | (-0.002, 0.012) |
| Constant | 0.793*** | 0.792*** | 0.800*** |
| | (0.786, 0.799) | (0.786, 0.799) | (0.793, 0.807) |
| Observations | 15,400 | 14,562 | 13,793 |
| $R^2$ | 0.00000 | 0.0001 | 0.0001 |
| Adjusted $R^2$ | -0.0001 | 0.00004 | 0.00004 |
| Residual Std. Error | 0.405 (df = 15398) | 0.405 (df = 14560) | 0.400 (df = 13791) |
| F Statistic | 0.015 (df = 1; 15398) | 1.622 (df = 1; 14560) | 1.608 (df = 1; 13791) |
| *Note:* | | | *p < 0.05. ** < 0.01, *** < 0.001 |

**Table S2.** Maternal Age and Spacing - population. Outcome is the difference between first- and secondborn children in sibship size two. If maternal age would need to be inversely associated with test scores, but they're positively associated with test scores. The effect of age on the difference between siblings is also much larger than mothers age.

| | *Dependent variable:* |
|---|---|
| | Diff. Between Sibs. |
| Age Difference Between Siblings | $-0.020^{***}$ |
| | (-0.022, -0.018) |
| Average Maternal Age At Birth | $0.002^{***}$ |
| | (0.002, 0.003) |
| Constant | $0.091^{***}$ |
| | (0.072, 0.110) |
| Observations | 406,270 |
| $R^2$ | 0.001 |
| Adjusted $R^2$ | 0.001 |
| Residual Std. Error | 0.873 (df = 406267) |
| F Statistic | $239.938^{***}$ (df = 2; 406267) |
| *Note:* | *p < 0.05. ** < 0.01, *** < 0.001 |

**Table S3.** Interaction between PGS and birth order. Child part of sample. Family-level fixed effects and between-families linear models.

**PGS x birth order interaction - children**

| | *Dependent variable:* | |
|---|---|---|
| | Z - Mean National Test Score | |
| | Family Fixed Effects OLS | OLS |
| | (1) | (2) |
| Birth Order | 0.002 | -0.060 |
| | (-0.145, 0.150) | (-0.131, 0.012) |
| PGS | 0.175*** | 0.219*** |
| | (0.089, 0.262) | (0.195, 0.243) |
| Female | 0.161 | 0.030 |
| | (-0.016, 0.338) | (-0.018, 0.078) |
| Sibship Size | | 0.005 |
| | | (-0.034, 0.044) |
| PC1 | | -0.827* |
| | | (-1.568, -0.087) |
| PC2 | | 1.138* |
| | | (0.196, 2.081) |
| PC3 | | -0.388 |
| | | (-1.596, 0.819) |
| PC4 | | 1.022 |
| | | (-0.465, 2.509) |
| PC5 | | -1.735* |
| | | (-3.295, -0.176) |
| PC6 | | -1.605* |
| | | (-3.069, -0.141) |
| PC7 | | 1.446 |
| | | (-0.006, 2.899) |
| PC8 | | 0.373 |
| | | (-1.105, 1.851) |
| PC9 | | -1.189 |
| | | (-2.698, 0.320) |
| PC10 | | 1.719* |

| | | |
|---|---|---|
| | | (0.393, 3.046) |
| Birth Order x PGS | -0.027 | 0.003 |
| | (-0.071, 0.017) | (-0.010, 0.017) |
| Birth Order x Female | -0.056 | -0.008 |
| | (-0.151, 0.039) | (-0.034, 0.019) |
| Birth Order x Sibship Size | | 0.005 |
| | | (-0.015, 0.026) |
| Constant | | 0.232*** |
| | | (0.105, 0.359) |
| Observations | 2,542 | 25,060 |
| $R^2$ | 0.728 | 0.079 |
| Adjusted $R^2$ | 0.448 | 0.078 |
| Residual Std. Error | 0.595 (df = 1251) | 0.787 (df = 25042) |
| F Statistic | | 125.823*** (df = 17; 25042) |
| *Note:* | | *p < 0.05. ** < 0.01, *** < 0.001 |

**Table S4.** Interaction between PGS and birth order. Parent part of sample. Family-level fixed effects and between-families linear models.

**PGS x birth order interaction - parents**

| | *Dependent variable:* | |
|---|---|---|
| | Z - Educational Attainment in Years | |
| | Family Fixed Effects OLS | OLS |
| | (1) | (2) |
| Birth Order | -0.166$^{***}$ | -0.080$^{***}$ |
| | (-0.254, -0.078) | (-0.109, -0.052) |
| PGS | 0.093$^{*}$ | 0.271$^{***}$ |
| | (0.021, 0.165) | (0.254, 0.287) |
| Female | 0.135$^{*}$ | 0.148$^{***}$ |
| | (0.007, 0.263) | (0.115, 0.181) |
| Sibship Size | | -0.053$^{***}$ |
| | | (-0.066, -0.039) |
| PC1 | | -0.104 |
| | | (-0.621, 0.414) |
| PC2 | | 1.475$^{***}$ |
| | | (0.805, 2.145) |
| PC3 | | -0.011 |
| | | (-0.881, 0.859) |
| PC4 | | -0.397 |
| | | (-1.468, 0.673) |
| PC5 | | -1.386$^{*}$ |
| | | (-2.500, -0.271) |
| PC6 | | -0.882 |
| | | (-1.959, 0.195) |
| PC7 | | 0.724 |
| | | (-0.346, 1.795) |
| PC8 | | -0.242 |
| | | (-1.326, 0.842) |
| PC9 | | -0.711 |
| | | (-1.805, 0.383) |
| PC10 | | -0.594 |
| | | (-1.577, 0.390) |

| | | |
|---|---|---|
| Birth Order x PGS | 0.029* | -0.002 |
| | (0.001, 0.057) | (-0.010, 0.005) |
| Birth Order x Female | 0.031 | 0.012 |
| | (-0.023, 0.085) | (-0.003, 0.027) |
| Birth Order x Sibship Size | | 0.006*** |
| | | (0.002, 0.009) |
| Constant | | 0.362*** |
| | | (0.297, 0.428) |
| Observations | 4,726 | 47,153 |
| $R^2$ | 0.745 | 0.098 |
| Adjusted $R^2$ | 0.420 | 0.097 |
| Residual Std. Error | 0.688 (df = 2075) | 0.881 (df = 47135) |
| F Statistic | | 299.604*** (df = 17; 47135) |
| *Note:* | | *p < 0.05. ** < 0.01, *** < 0.001 |

**Table S5.** Parental educational attainment and average difference between first- and secondborn in sibship size 2 - Sample. Outcome is the difference between first- and secondborn. Any positive coefficient from parental educational attainment indicate that higher educated parents have siblings where the difference between the first- and the secondborn is larger than those with lower education.

**Difference between birth order 1 and 2 - sample**

| | *Dependent variable:* | | | | | | |
|---|---|---|---|---|---|---|---|
| | Difference Between 1st and 2nd Born | | | | | | |
| | OLS | OLS | OLS | OLS | OLS | OLS | OLS |
| | (1) | (2) | (3) | (4) | (5) | (6) | (7) |
| Z - Parental EA in Years | 0.063** | 0.046 | 0.051 | | 0.046 | 0.051 | |
| | (0.018, 0.108) | (-0.004, 0.096) | (-0.002, 0.103) | | (-0.004, 0.096) | (-0.002, 0.103) | |
| Parental EA - Medium | | | | 0.147 | | | 0.111 |
| | | | | (-0.069, 0.364) | | | (-0.120, 0.343) |
| Parental EA - High | | | | 0.232* | | | 0.206 |
| | | | | (0.026, 0.438) | | | (-0.014, 0.426) |
| PGS | | 0.037 | 0.033 | 0.038* | 0.037 | 0.033 | 0.035 |
| | | (-0.0002, 0.075) | (-0.006, 0.073) | (0.001, 0.075) | (-0.0002, 0.075) | (-0.006, 0.073) | (-0.004, 0.073) |
| Female | | 0.071 | 0.058 | 0.074 | 0.071 | 0.058 | 0.060 |
| | | (-0.006, 0.147) | (-0.024, 0.140) | (-0.003, 0.150) | (-0.006, 0.147) | (-0.024, 0.140) | (-0.022, 0.142) |
| Birth Length | | | -0.002 | | | -0.002 | -0.004 |
| | | | (-0.032, 0.029) | | | (-0.032, 0.029) | (-0.035, 0.027) |
| Gestational Age | | | 0.008 | | | 0.008 | 0.008 |
| | | | (-0.014, 0.030) | | | (-0.014, 0.030) | (-0.014, 0.030) |
| Birth Weight | | | -0.00003 | | | -0.00003 | -0.00003 |
| | | | (-0.0002, 0.0001) | | | (-0.0002, 0.0001) | (-0.0002, 0.0001) |
| Constant | 0.037 | -0.061 | -0.156 | -0.231 | -0.061 | -0.156 | -0.227 |

| | | | | | | | |
|---|---|---|---|---|---|---|---|
| | (-0.016, 0.089) | (-0.187, 0.066) | (-1.511, 1.198) | (-0.466, 0.004) | (-0.187, 0.066) | (-1.511, 1.198) | (-1.583, 1.129) |
| Observations | 2,161 | 1,929 | 1,711 | 1,929 | 1,929 | 1,711 | 1,711 |
| $R^2$ | 0.003 | 0.007 | 0.007 | 0.009 | 0.007 | 0.007 | 0.008 |
| Adjusted $R^2$ | 0.003 | 0.005 | 0.003 | 0.006 | 0.005 | 0.003 | 0.004 |
| Residual Std. Error | 0.844 (df = 2159) | 0.855 (df = 1925) | 0.847 (df = 1704) | 0.854 (df = 1924) | 0.855 (df = 1925) | 0.847 (df = 1704) | 0.846 (df = 1703) |
| F Statistic | 7.488** (df = 1; 2159) | 4.199** (df = 3; 1925) | 1.909 (df = 6; 1704) | 4.141** (df = 4; 1924) | 4.199** (df = 3; 1925) | 1.909 (df = 6; 1704) | 2.023* (df = 7; 1703) |
| *Note:* | | | | | | | *p**p***p<0.001 |

**Table S6.** Parental and Sibling Genetic Nurture. Child part of the sample.

**Parental and sibling genetic nurture**

| | *Dependent variable:* | | | | | |
|---|---|---|---|---|---|---|
| | Z - Mean National Test Score | | | | | |
| | (1) | (2) | (3) | (4) | (5) | (6) |
| Own PGS | 0.203*** | 0.187*** | 0.193*** | 0.171*** | 0.187*** | 0.175*** |
| | (0.175, 0.232) | (0.152, 0.221) | (0.158, 0.228) | (0.127, 0.214) | (0.153, 0.221) | (0.131, 0.219) |
| Female | 0.046 | 0.049 | 0.039 | 0.040 | 0.046 | 0.041 |
| | (-0.011, 0.102) | (-0.010, 0.107) | (-0.021, 0.098) | (-0.020, 0.101) | (-0.012, 0.103) | (-0.020, 0.103) |
| Mothers PGS | | 0.027 | | 0.029 | | 0.011 |
| | | (-0.009, 0.062) | | (-0.009, 0.067) | | (-0.033, 0.054) |
| Fathers PGS | | | 0.021 | 0.030 | | 0.015 |
| | | | (-0.015, 0.057) | (-0.008, 0.068) | | (-0.029, 0.058) |
| Sibling PGS | | | | | 0.045** | 0.037 |
| | | | | | (0.011, 0.079) | (-0.007, 0.081) |
| Constant | 0.162*** | 0.152** | 0.172*** | 0.162*** | 0.163*** | 0.158** |
| | (0.074, 0.251) | (0.061, 0.243) | (0.079, 0.265) | (0.067, 0.257) | (0.073, 0.252) | (0.062, 0.255) |
| Observations | 2,861 | 2,739 | 2,605 | 2,518 | 2,762 | 2,436 |
| $R^2$ | 0.064 | 0.063 | 0.064 | 0.064 | 0.071 | 0.069 |
| Adjusted $R^2$ | 0.063 | 0.062 | 0.063 | 0.062 | 0.070 | 0.067 |
| Residual Std. Error | 0.773 (df = 2858) | 0.775 (df = 2735) | 0.776 (df = 2601) | 0.776 (df = 2513) | 0.770 (df = 2758) | 0.774 (df = 2430) |
| F Statistic | 97.878*** (df = 2; 2858) | 61.365*** (df = 3; 2735) | 59.590*** (df = 3; 2601) | 42.671*** (df = 4; 2513) | 70.583*** (df = 3; 2758) | 36.018*** (df = 5; 2430) |
| *Note:* | | | | | | *p < 0.05. ** < 0.01, *** < 0.001 |

### SI References

1. M. J. Knol, W. R. Pestman, D. E. Grobbee, The (mis)use of overlap of confidence intervals to assess effect modification. *Eur. J. Epidemiol.* **26**, 253–254 (2011).
